# Machine learning approaches for the identification and analysis of enterotoxin genes in *Staphylococcus aureus* genomes

**DOI:** 10.64898/2026.05.01.722155

**Authors:** Alex Uttin, Richard M. Leggett, Vincent Moulton, Jo Dicks

**Author notes:** **Corresponding author and email address** Jo Dicks.

## Abstract

2.

*Staphylococcus aureus* produces a broad range of enterotoxins that act as superantigens, disrupting host immune responses and resulting in a myriad of clinical symptoms. However, large-scale analyses determining enterotoxin gene diversity, lineage structure and isolate metadata remain scarce. We analysed 15,887 *S. aureus* RefSeq genomes using a machine learning pipeline combining profile Hidden Markov Model-based enterotoxin gene identification, lineage typing, gene profile-based strain clustering and association rule mining using a broad range of gene and metadata features. This approach identified 35 distinct enterotoxin genes and five variant forms, including two putative novel enterotoxin genes, *sel34* and *sel35*. HDBSCAN clustering distinguished 45 enterotoxin gene profile groups, revealing strong associations between the two major egc enterotoxin gene cluster variants (OMIWNG and OMIUNG) and Clonal Complex membership: *CC5, CC22* and *CC45* with OMIWNG; *CC30* and *CC121* with OMIUNG. Integration of isolate metadata exposed distinct geographic and temporal trends, including a recent rise in non-egc lineages derived from Asia and animal sources. These findings show that *S. aureus* enterotoxin diversity is structured by lineage, mobile genetic element composition and Clonal Complex association. The discovery of *sel34* and *sel35*, together with the comprehensive overview of lineage-specific enterotoxin profiles, expands current understanding of *S. aureus* virulence evolution and provides a scalable analytical framework for monitoring toxin gene dynamics in clinical and environmental populations.

**Impact Statement:** Understanding how virulence genes evolve and spread in *Staphylococcus aureus* is vital for predicting pathological potential and managing infection risk for this species. By analysis of over 15,000 publicly available *S. aureus* genomes, this study provides the most comprehensive overview to date of enterotoxin gene diversity and lineage structure. Using machine learning and large-scale genomic mining, we reveal clear evolutionary and epidemiological patterns linking enterotoxin gene clusters to specific Clonal Complexes and identify two previously unknown enterotoxin genes. These findings highlight how recombination and horizontal gene transfer shape *S. aureus* toxins across hosts, continents and time. The resulting analytical framework offers a scalable foundation for future genomic surveillance of virulence evolution in both clinical and environmental settings.

**Data Summary:** Genome assemblies and associated metadata for the 15,887 *Staphylococcus aureus* strains analysed within this study were generated elsewhere prior to this study and were here downloaded from the NCBI RefSeq database. Secondary datasets and descriptions of algorithmic approaches used to develop them are presented in the main text and within the Supplementary Files 1 and 2.

## 5. Introduction

*Staphylococcus aureus* is a Gram-positive, spherical bacterium that typically forms grape-like clusters. As an opportunistic pathogen, it causes a range of infections in humans, from mild skin conditions to severe diseases such as pneumonia, toxic shock syndrome and sepsis (1). Approximately 20% of the world’s population are believed to be persistent carriers of *S. aureus*, 60% are intermittent carriers and 20% never harbour the organism (2). The management and treatment of *S. aureus* infections are increasingly complicated by the emergence of antibiotic-resistant strains, notably Methicillin-resistant *Staphylococcus aureus* (MRSA) and multidrug-resistant strains. Methicillin-susceptible *S. aureus* (MSSA) can acquire antibiotic resistance genes via horizontal gene transfer mediated by mobile genetic elements, thereby converting to MRSA (3). In 2007, MRSA infections were estimated to cost the UK National Health Service approximately £1 billion annually (4), a figure that has likely risen since that time. Across Europe, MRSA is projected to affect over 150,000 patients annually, underscoring the urgent need for innovative strategies to mitigate its impact on hospital capacity and healthcare costs (5).

*S. aureus* exerts its pathogenic effects partly through the production of superantigens namely, enterotoxins and enterotoxin-like proteins plus the toxic shock syndrome toxin-1 (6). Unlike conventional antigens that typically stimulate around 0.001% of T cells, these superantigens can activate up to 20% of T cells, triggering a cytokine storm that may result in severe clinical manifestations, including seizures (7). The nomenclature for the enterotoxin and enterotoxin-like genes follows the system proposed by Lina *et al*., 2004 and continued by Fischer *et al*., 2019 where emetic-confirmed enterotoxin genes are prefixed with “se” and other enterotoxin-like genes with “sel” (8,9). To date 33 enterotoxin and enterotoxin-like genes (*sea, seb, sec1, sed, see, seg, seh, sei, selj, sek, sel, sem, sen, seo, sep, seq, ser, ses, set, selu, selv, selw, selx, sely, selz, sel26, sel27, sel28, sel29p, sel30, sel31, sel32, sel33*), in addition to 5 variant forms (*sec2, sec3, seh-2p, ses-2p, ses-3p*), have been identified and found to be distributed throughout the *S. aureus* genome. Perhaps unsurprisingly for such a large gene family, paralogues often cluster together; the most prominent of these gene clusters is the enterotoxin gene cluster (egc), which primarily exists in two variant forms of six genes from a repertoire of nine (10). Specific enterotoxins/superantigens are causal to individual conditions; *tsst-1* causes toxic shock syndrome and *seb* has been classified as a bio-terrorist threat (11,12) though many are not associated currently with an exclusive pathology.

Despite advances in our understanding of *S. aureus* prevalence and transmission, a comprehensive overview that integrates enterotoxin gene profiles, lineage information and source metadata remain lacking. Dicks *et al*., 2021 began such an approach with their analysis of 133 NCTC *S. aureus* isolates and a brief exploration of larger public datasets (10). In the present study, we expanded upon their methodology using machine learning approaches by analysing 15,887 RefSeq *S. aureus* isolates, thereby incorporating a broader exploration of the bacterium’s pathology and epidemiology. Our initial objective was to investigate the associations between enterotoxin gene counts and identities across different *S. aureus* lineages. Ultimately, our study has revealed novel genes, distinct enterotoxin groupings, and previously unrecognised relationships between lineages and enterotoxins, findings that were further enriched by the integration of extensive *S. aureus* metadata.

## 6. Methods

### RefSeq Dataset Preparation

The genome assemblies of 15,887 *Staphylococcus aureus* isolates were downloaded from The National Center for Biotechnology Information (NCBI) RefSeq database (https://www.ncbi.nlm.nih.gov/refseq/) on 30/11/2023. In order to identify the DNA sequences of putative enterotoxin genes within the RefSeq assemblies, HMMER 3.4 (13,14) was used in combination with two profile Hidden Markov Models (pHMMs) developed within a previous study of Staphylococcal enterotoxins (10). Two conditions were used to filter the results of the pHMM search: a ‘mid-filter’ which required an E-value of 1.0e-10 or less and enterotoxin-like sequences of near complete length (i.e. covering at least positions 4 to 601 of the HMM for *selx* and *tsst-1* and positions 252 to 753 for other enterotoxin genes) and a ‘no-filter’ variant requiring only an E-value of 1.0e-10 or less. HMMER produced a FASTA file for each filtered dataset containing the putative enterotoxin gene sequence attached to the name of the parent shotgun sequence.

Dicks *et al*.’s earlier study used the sequences of 27 target enterotoxin genes obtained from UniProt (https://www.uniprot.org/) and GenBank (https://www.ncbi.nlm.nih.gov/genbank/) around which to cluster sequences identified within their strains of interest, expanding the list of enterotoxin gene forms to 38, comprising 33 genes and 5 gene variants (10). These 38 reference sequences plus that of the closely related gene *tsst-1* were integrated into the appropriate FASTA file for subsequent alignment with MAFFT (version 7, https://mafft.cbrc.jp/alignment/software/), employing the strategy: *G-large-INS-1*, scoring matrix: *20PAM/k=2* parameter and a high gap opening penalty (15–17). This process returned four FASTA files of the putative enterotoxin genes within the 15,887 genome assemblies (i.e. s*elx*/*tsst-1* and other enterotoxin genes for each of the two filter conditions) now grouped around the reference enterotoxin genes.

The multiple sequence alignment software AliView (version 1.3, https://ormbunkar.se/aliview/) enabled manual slicing of the aligned sequences into distinct groups that either centred around a reference enterotoxin gene or formed a new group with no reference gene (18). As putative enterotoxin genes within the alignment were named according to the shotgun sequences in which they were found, rather than the genome assembly, Python dictionaries linking genome assembly identifiers to shotgun sequence identifiers and shotgun sequence identifiers to reference sequence identifiers were created. This enabled a table of whole genome assemblies and the putative enterotoxin genes contained within them to be generated.

The mlst software (version 2.23.0, https://github.com/tseemann/mlst) was employed to determine the allelic forms of schema “housekeeping” genes for each RefSeq genome assembly. Each strain’s allelic combination was then input to the online BIGSdb tool to generate the appropriate lineage designation for each individual genome assembly (19,20). The lineage data was subsequently combined with the enterotoxins found within each isolate, resulting in a table containing: Strain name, enterotoxin gene presence/absence, Sequence Type and Clonal Complex. This table is henceforth referred to as the RefSeq dataset.

### Clustering of enterotoxin gene profiles

The binary enterotoxin gene data in the RefSeq dataset was extracted for use in cluster analysis. First, the dimensionality reduction algorithm t-SNE was used to produce a 2D representation of the enterotoxin gene data that preserved the similarities of the gene presence/absence data between isolates (21). Sixty-four different t-SNE variations were produced with varied parameter values and finalised as: *Components: 2, Perplexity: 100, Iter: 2000* and *Learning rate: 200*, selected visually based on expected cluster shape and density. This t-SNE model served as a visual aid for inspection of the subsequent generated clusters (e.g. see Figure S1 in Supplementary File 1).

Second, the density-based clustering method DBSCAN (scikit-learn version 1.7.1, https://scikit-learn.org/stable/modules/generated/sklearn.cluster.DBSCAN.html) was used to group the enterotoxin gene data. DBSCAN was chosen initially due to its ability to distinguish clusters from noise in high dimensional data and arbitrarily shaped clusters (22). However, DBSCAN has limitations when dealing with datasets containing clusters of varying densities, which we suspected might be the case with the RefSeq dataset. Subsequent analyses were therefore conducted with the HDBSCAN algorithm (23,24). HDBSCAN extends DBSCAN by building a hierarchy of clusters using a range of density thresholds, identifying sub-clusters and collapsing this hierarchy to extract the most stable and meaningful clusters.

576 HDBSCAN models were produced with the parameters: *Minimum cluster size (5, 10, 15, 20, 40, 80), Minimum sample size (5, 10, 15, 20, 40 80), epsilon value (0.1, 0.3, 0.5, 0.7), metric choice (euclidean, manhattan)* and *cluster selection metric (eom, leaf)*. The HDBSCAN models were applied to the pre-selected t-SNE model to produce 576 scatter graphs for visual analysis and comparison. Four final models were proposed using visual analysis and cluster count from the 576 HDBSCAN variations leading to a final visually judged HDBSCAN model with the parameters: *Minimum cluster size: 40, Minimum sample size: 20, Epsilon value: 0.3, Metric: manhattan* and *Cluster selection metric: eom*. The relevant cluster label produced by the HDBSCAN model was appended to each isolate in the RefSeq dataset table. The HDBSCAN model consisted of 44 clusters plus a noise cluster with an average strain count of 331, a standard deviation of 142 with a high of 2,821 isolates and a low of 41 isolates, the limit set by the minimum sample size parameter. The 975 noise cluster isolates (label: “-1”) were sub-clustered by adjusting parameters to *Minimum cluster size: 10* and *Minimum sample size: 10* to account for the large size of the RefSeq noise cluster.

### Isolate metadata and association rule mining

The NCBI genome dataset includes the option to download GenBank Sequence and annotation files (GBFF), which can provide additional information for each isolate. The GBFF files can include strain isolation date, isolation country and isolation host. Of the 15,887 RefSeq dataset isolates, 14,077 (88.6%) contained isolation countries, 13,386 (84.3%) contained isolation years and 11,700 (73.6%) contained isolation host. The RefSeq dataset was finalised to include Isolate ID, enterotoxin gene presence/absence, Sequence Type, Clonal Complex, HDBSCAN Cluster, isolation metadata information and isolation publication information. The process of creating the RefSeq dataset, complete with metadata, is visualised in Figure 1.

**Figure 1:**
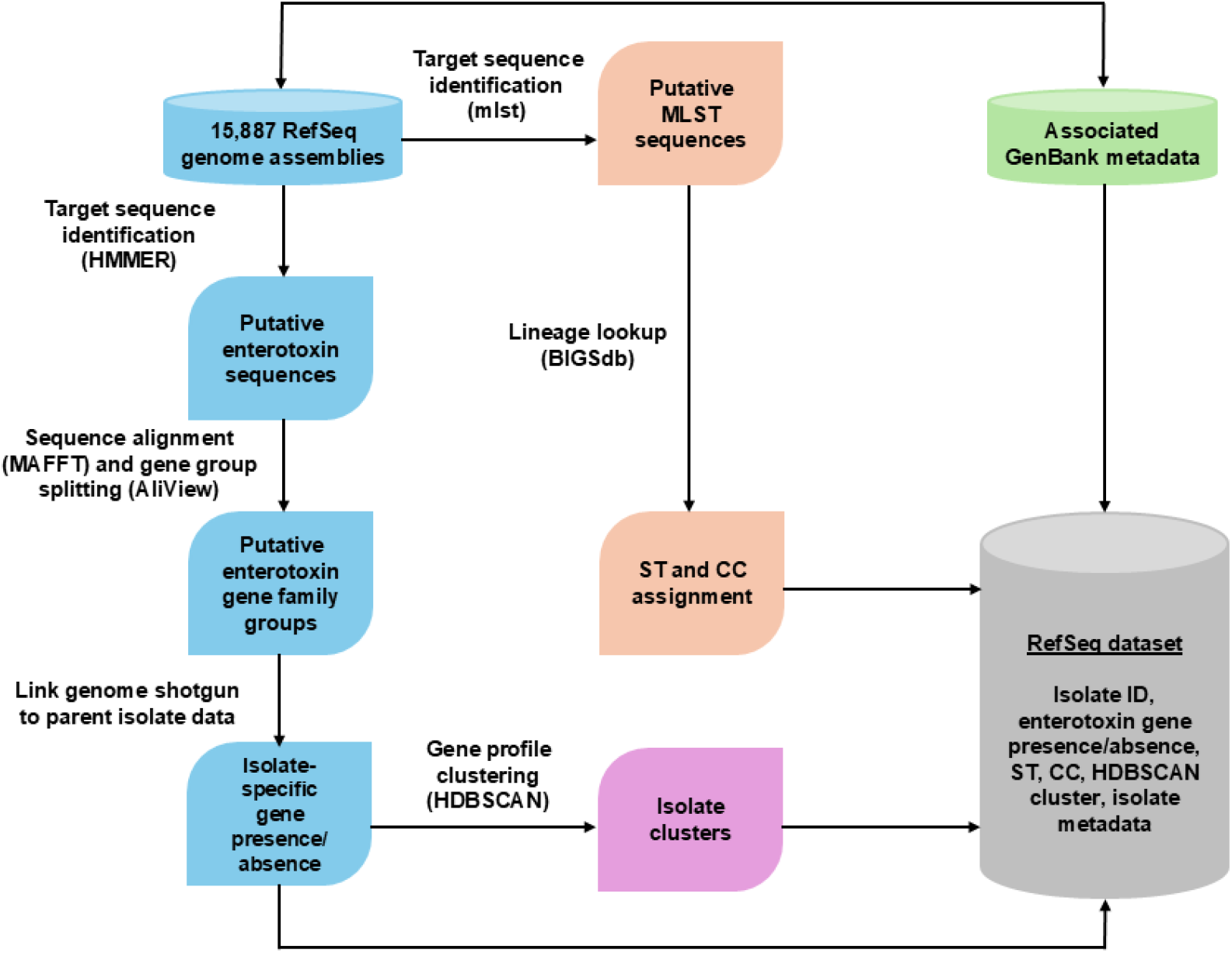
A pipeline for generating the RefSeq dataset for 15,887 isolates downloaded from the RefSeq database. The individual pipeline processes are colour-coded for clarity. The blue pipeline follows the process to identify putative enterotoxin genes present in each genome assembly. The orange pipeline determines the lineage information for each genome assembly. The green pipeline represents the inclusion of isolate metadata from GenBank and the purple pipeline the HDBSCAN clustering of the isolates based on their enterotoxin gene profiles.

The apriori association rule mining algorithm was used to generate rules between enterotoxin genes, enterotoxin genes to lineage and enterotoxin genes to metadata (25,26). The rule mining algorithm did not initially provide meaningful results when attempting to associate enterotoxin genes with lineage and metadata. The enterotoxin genes *sel26* and *selx* were subsequently removed from the dataset used for rule mining due to their high prevalence across the RefSeq dataset, a likely product of their chromosomal location. Furthermore, the enterotoxin genes that comprise the egc enterotoxin gene cluster were abstracted into their respective egc variant (OMI(U/W)NG), source countries were abstracted to source continents, years to time periods and hosts to animal or human. The apriori algorithm, acquired through the MLxtend Python package, was then deployed using the parameters: *Minimum support: 0.005, Minimum threshold: 0.6* and *Metric: confidence* (27). Lower support and threshold values were used to ensure smaller patterns within the RefSeq dataset were not ignored.

Networks were generated from the apriori association mined rules using the NetworkX Python package (28). To simplify the networks, the association rules were unidirectionally flattened from antecedents to consequents by averaging the confidence values for each unique pair of associated enterotoxin genes, Clonal Complex or isolation metadata and normalising all results to create a strength score between 0 and 1. This process is described further in Sections 2 and 3 of Supplementary File 1.

### Phylogenetic and recombinant sequence analysis

Representative amino acid sequences of groups that did not cluster around a reference enterotoxin gene were aligned manually using AliView to the amino acid multiple sequence alignment used in Dicks *et al*., 2021 (10,18). An amino acid phylogenetic tree was then estimated for the enterotoxin gene sequences using this new alignment as input to IQ-TREE with 1000 bootstraps, following model selection with ModelFinder (29). The enterotoxin gene *sel33*, (a recombinant of *selw* and *sen*) was excluded from the analysis as its presence can distort the resulting topology. Graphical representation and re-rooting of the phylogeny were carried out with FigTree (http://tree.bio.ed.ac.uk/software/figtree/).

A Neighbor-Net was generated from the alignment with *sel33* present using SplitsTree v6.3.34 (30,31). A nucleotide multiple sequence alignment was hand-aligned from the amino acid alignment. The amino acid sequences of the two *ses* variants, *ses-2p* and *ses-3p*, were created from fragmented sequences. As a result, identifying a suitable nucleotide sequence proved challenging and *ses-2p* and *ses-3p* were excluded from the enterotoxin nucleotide alignment. This resulting nucleotide alignment was used as input to RDP5 in order to identify putative recombinant enterotoxin gene sequences (32).

## 7. Results

### The RefSeq dataset contains two novel putative enterotoxin-like genes, *sel34* and *sel35*

Analysis of the RefSeq dataset established the enterotoxin gene composition across the 15,887 *S. aureus* isolates, with 35 distinct enterotoxin genes plus 5 variant forms of 4 of the genes (see the RefSeqDataset worksheet within Supplementary File 2). Among this dataset, two distinct novel putative enterotoxin-like gene sequences were identified and named *sel34* and *sel35* (see Section 5 of Supplementary File 1), according to nomenclature established by Lina *et al*., 2004 and used subsequently in numerous studies (8,9). The two novel enterotoxin genes were present in both the mid-filter and no-filter pHMM searches. The two isolates in the RefSeq dataset that harbour these genes were collected in 2000 and 2013 from Kenya and Somalia respectively, with both putative enterotoxin genes identified in each isolate. The genome of one isolate contained only the *sel34* and *sel35* enterotoxin genes, while the other contained *ser, selj, ses, set, sel34* and *sel35*. Neither isolate contained the most prevalent enterotoxin genes, *sel26* and *selx*. The isolates were found to belong to Sequence Types 3602 and 3576, however the BIGSdb software at the PubMLST site was unable to find a Clonal Complex designation for these two Sequence Types.

Querying the nucleotide sequences of *sel34* and *sel35* in GenBank provided three high-scoring hits for each gene. All hits originated from strains isolated from East African Camels from the Horn of Africa within a study by Akarsu *et al*. (33). A phylogenetic tree of the amino acid sequences of 39 of the 40 enterotoxin sequences plus *tsst-1* was estimated using IQ-TREE, as shown in Figure 2. The various clades seen within Figure 2 were largely consistent with those identified in a recent study (10); the amino acid sequence of *sel34* was found to most closely resemble that of *seb* with 67% similarity, while *sel35* grouped within the clade containing genes such as *sel26* and *sea*, exhibiting a 65% similarity to *sel26*.

**Figure 2:**
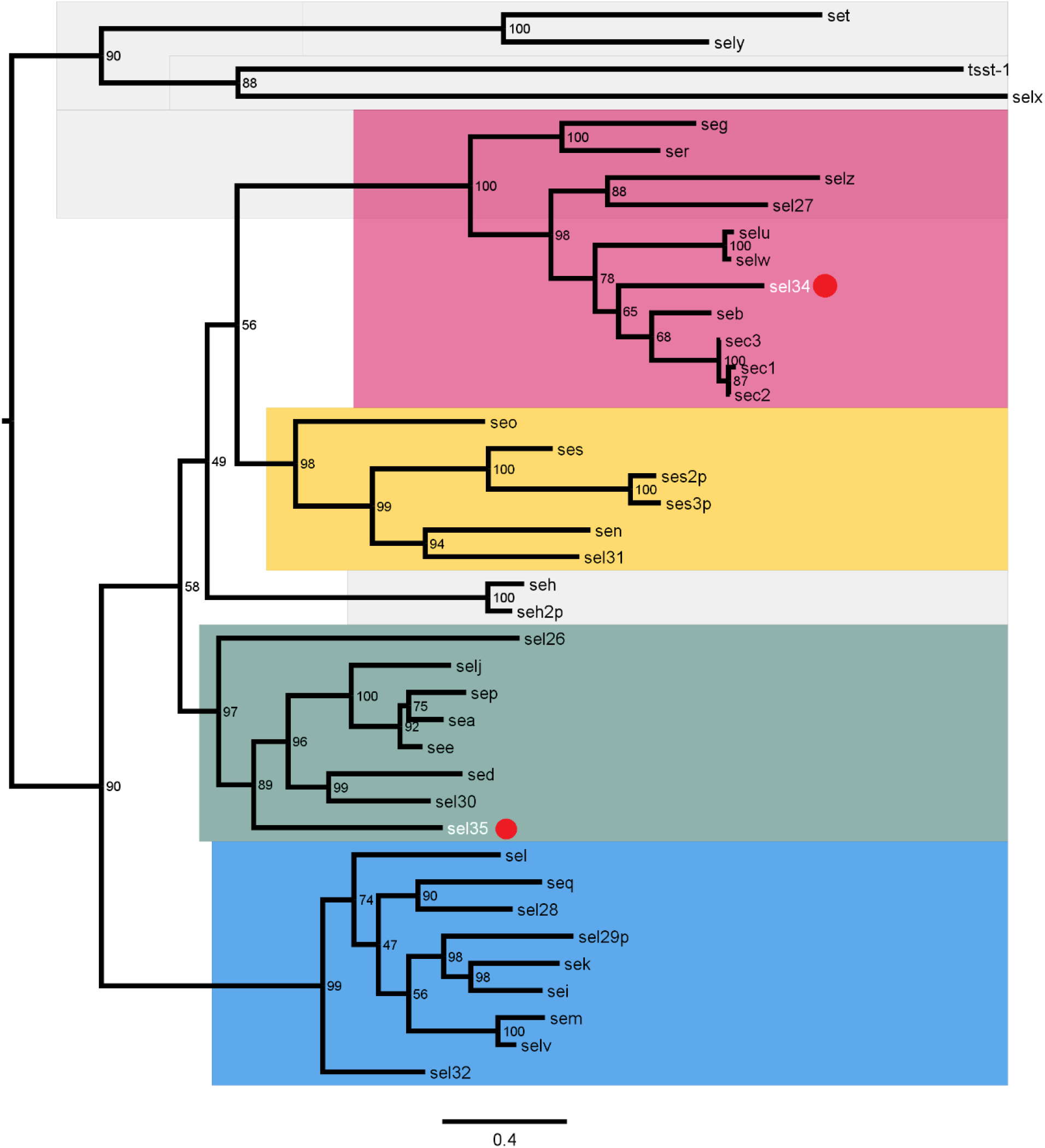
A phylogenetic tree of the 40 enterotoxin genes/gene variants in the RefSeq dataset minus *sel33*. The amino acid phylogeny of *Staphylococcus aureus* enterotoxin genes found in the RefSeq dataset. *Sel33* is not shown as the gene’s recombinant origin distorts the topology of the phylogeny. Compact gene groups (clades) are highlighted in coloured blocks. The phylogenetic tree was generated using IQ-TREE with the VT+F+R4 model. One thousand bootstrap trees were performed, with the percentage of supporting bootstraps for each split displayed in the internal node. The tree was edited, coloured and re-rooted in FigTree with two novel enterotoxin-like sequences (*sel34* and *sel35*) highlighted with white text and a red circle.

### Recombination has played a role in the evolution of the enterotoxin gene family

Analysis of the 40 enterotoxin nucleotide sequences with the RDP5 software suggested 22 potential recombinant enterotoxin genes, 10 with one unknown major parent (from whom the majority of the sequence under analysis was inherited/obtained), 7 with unknown minor parents (donor of a smaller section of the sequence), 4 putative recombinants with two known parents and 1 duplicate recombinant. The 4 putative recombinants with known parents – the most well supported class of results - were *sel33, selv, seh-2p* and *sel34*. RDP hypothesised *sel33* to be a recombinant of *selw* and *sen, selv* a recombinant of both *sem-sei* and *sel28*-*sel27, seh-2p* a recombinant of *seh and selw* and *sel34* a recombinant of *seb* and *sely*.

It has already been demonstrated that *selv* and *sel33* are both likely recombinants of neighbouring *egc* gene cluster genes (10,34), involving *sem*/*sei* and *selw*/*sen* respectively, giving weight to these results. While the recombinant status of *seh-2p* may have been affected by the manual sequence reconstruction process required for sequence alignment, the indication that *sel34* might also be a recombinant gene warrants further investigation, as do the additional results potentially involving as yet unidentified gene family members.

A NeighborNet was estimated using the amino acid sequences of all 40 enterotoxin genes/gene variants plus *tsst-1* using SplitsTree, as shown in Figure 3. The NeighborNet represents a complex split system indicating that departures from treelike evolutionary processes, such as recombination events, may have taken place. Notably, two of the more distant splits (e.g. potentially indicating the most recent events) involve *selv* and *sel33*. However, support of *sel34*’s potential recombinant origin was not clearly apparent.

**Figure 3:**
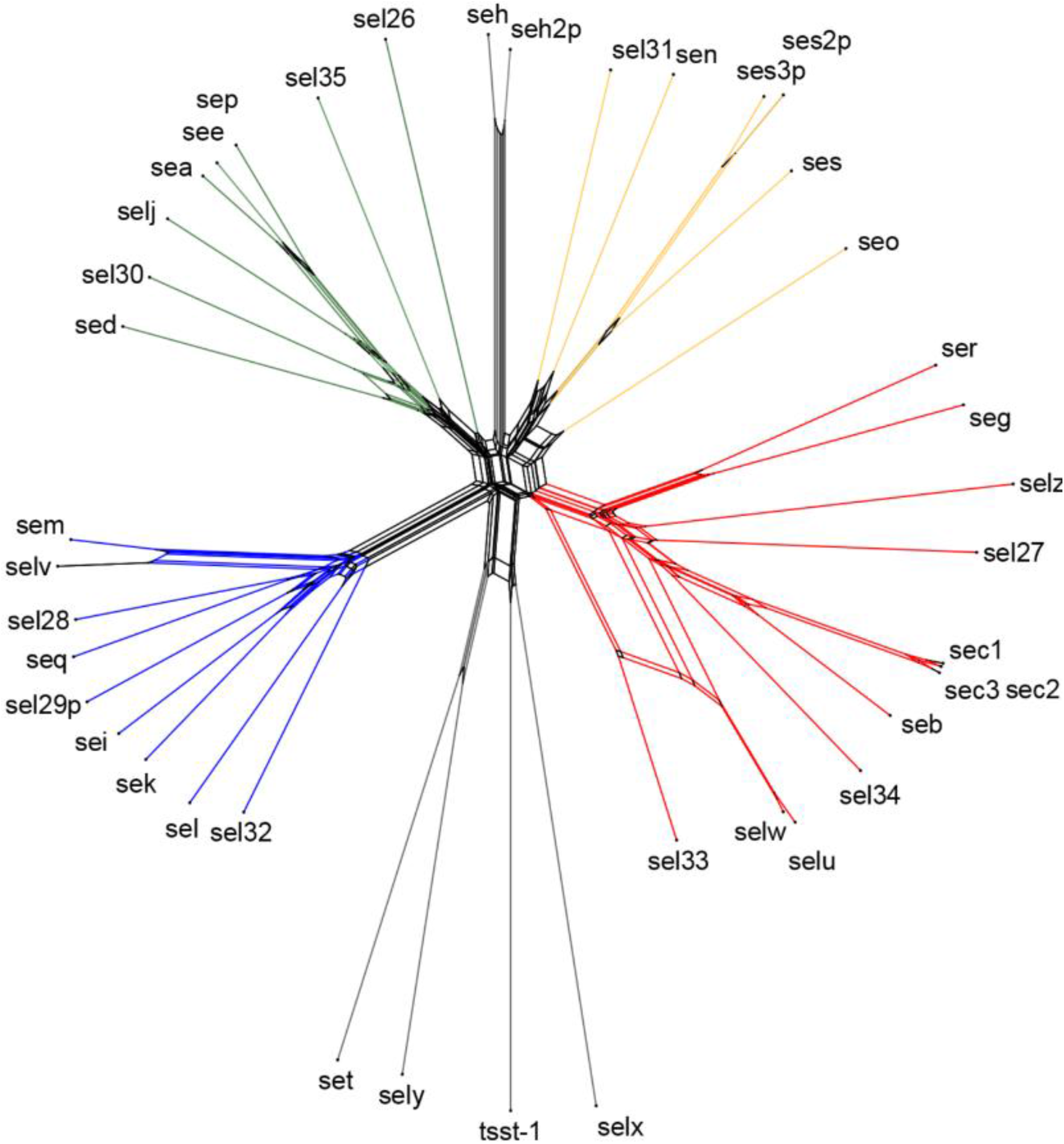
A NeighborNet of all 40 *Staphylococcus aureus* enterotoxin genes and gene variants found within the RefSeq dataset, plus the closely related *tsst-1* gene. The NeighborNet was generated using SplitsTree v6.3.34 from the 41 amino acid sequences which included the newly discovered enterotoxin gene family members *sel34* and *sel35*. The NeighborNet is coloured with the same clades found in Figure 2.

### The prevalence of enterotoxin genes and Clonal Complex membership within the RefSeq dataset varies considerably

Together two enterotoxin genes, *sel26* and *selx*, were present across more than 80% of the 15,887 isolates of the RefSeq dataset with *sel26* appearing in 96% of isolates followed by *selx* in 89% of isolates. Six other enterotoxin genes, *seo, sem, sei, selw, sen* and *seg* (collectively forming a key variant of the egc enterotoxin gene cluster: OMIWNG) were found in more than 40% of the *S. aureus* isolates, with individual gene frequencies ranging from 49% down to 43%. Despite being known as a key member of the egc gene cluster, *selu* was found in only 6% of isolates. The *tsst-1* gene, causal agent of toxic shock syndrome, and the *seb* gene, linked to immune system collapse and therefore considered a bioterrorist threat, both appeared in 7% of *S. aureus* isolates within the RefSeq dataset.

Application of the mlst software to the RefSeq dataset reported 667 Sequence Types and 10 Clonal Complexes, with 220 isolates lacking an associated Sequence Type and 3,178 (20%) without a Clonal Complex designation. Of the Clonal Complexes, *CC5* was most common with 4,414 appearances (28%), followed by *CC8* at 3,611 appearances (23%). *CC121* contained the highest average count of enterotoxin genes at 10 enterotoxin genes for each isolate and *CC15* had the lowest average count with 2 enterotoxin genes. The most populous Clonal Complexes *CC5* and *CC8* had significantly different counts of enterotoxin genes at 38,712 and 14,068 respectively, displaying the vast differences in enterotoxin gene count between lineages.

Of the 667 Sequence Types, 376 were observed only once and 583 Sequence Types were found in fewer than ten *S. aureus* isolates each. ST5, the canonical strain of *CC5*, was unsurprisingly the most frequently observed Sequence Type with 3,044 appearances, at 69% of *CC5* and 19% of the RefSeq dataset, followed by *ST8* at 2,527 appearances.

### Prevalence of egc gene cluster forms differ between Clonal Complexes and in metadata classes

The egc enterotoxin gene cluster typically consists of six enterotoxin genes: *seo, sem, sei, selu/selw, sen* and *seg*. The cluster type possessed by most individual strains can therefore be referred to as either OMIUNG or OMIWNG, indicating the physical order of the egc genes and the presence of either *selu* or *selw*. Additional egc gene cluster variants such as OMIUN and OVUNG also exist but are believed to be rare. In the RefSeq dataset, approximately 6,776 OMIWNG egc gene cluster variants and 1,000 OMIUNG variants were observed (using presence of *selu*/*selw* as a proxy, see the RefSeqCC worksheet in Supplementary File 2). A clear Clonal Complex lineage split was observed between OMIUNG and OMIWNG, along with distinct differences in metadata. For example, while *CC5* isolates were found to contain approximately 3,939 copies of the OMIWNG egc enterotoxin gene cluster (89% of *CC5* isolates and 25% of all isolates), those in *CC8* contained just 164 OMIWNG egc enterotoxin gene cluster variants (5% of *CC8* isolates and 1% of all isolates). This observation is further supported by Figure 4 where egc gene cluster presence/absence can be observed across all Clonal Complexes, with all but *CC1* showing a clear preference of gene cluster type.

**Figure 4:**
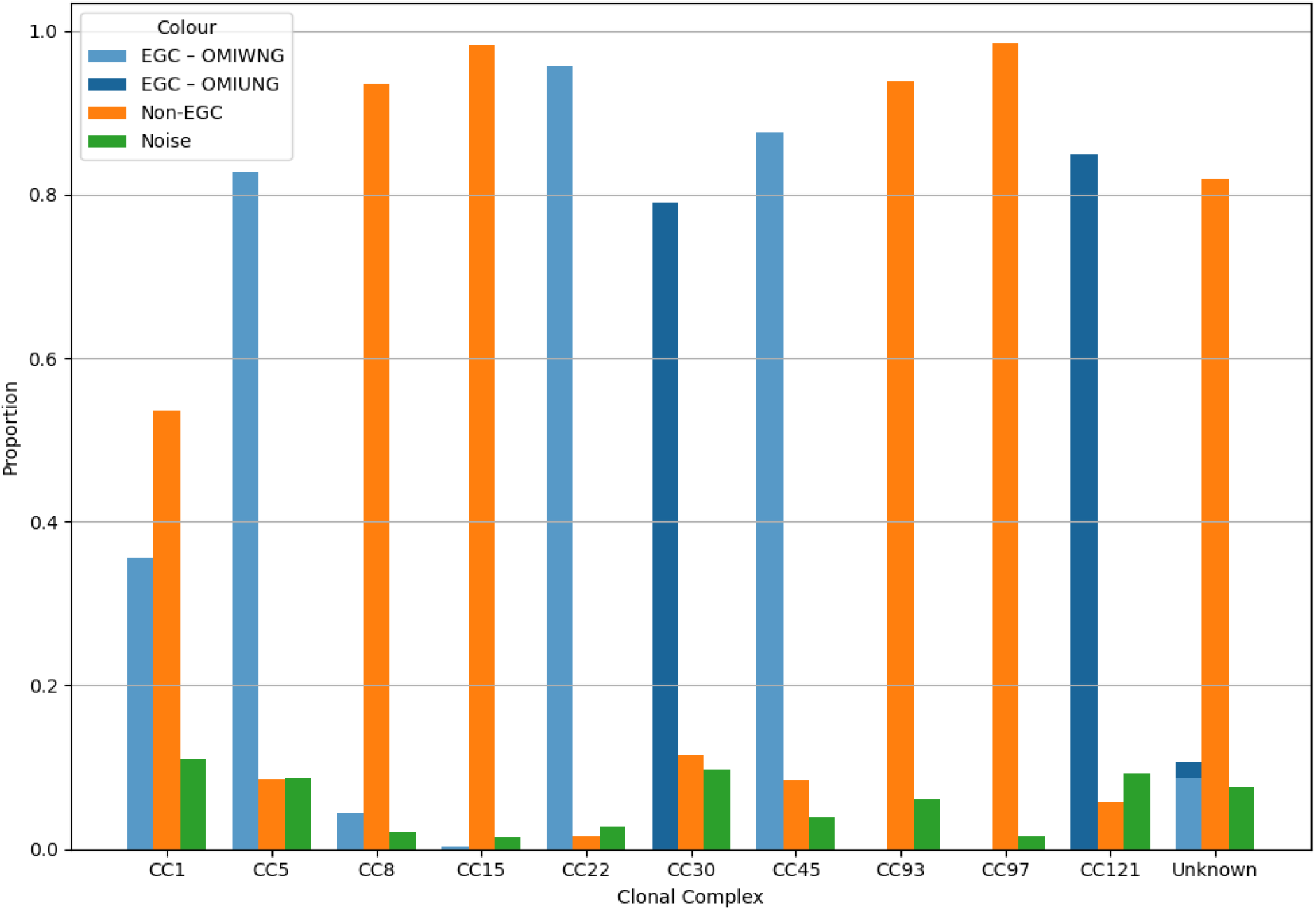
HDBSCAN cluster composition of the strains belonging to the ten identified Clonal Complexes (with remaining strains assigned to an Unknown group). Blue bars indicate clusters comprised of strains whose genomes contained an egc gene cluster and have been shaded according to the egc gene cluster variant, orange for clusters comprised of strains lacking an egc gene cluster (Non-EGCC) and green for strains within the noise cluster. Most Clonal Complexes are predominantly comprised of strains from one type of HDBSCAN cluster only, with the notable exception of *CC1*.

Overall, the OMIWNG variant was found in 43% of RefSeq genomes, primarily concentrated in *CC5, CC22*, and *CC45* isolates (see the RefSeqCC worksheet in Supplementary File 2). OMIWNG showed a strong negative Pearson correlation coefficient with *CC8* (-0.42) along with more moderate negative correlations with *CC30* (-0.18) and Unknown Clonal Complexes (-0.33). OMIWNG showed a high prevalence in isolates from North and South America (56% and 48% respectively), but much lower frequencies in isolates from Africa (18%; see the ContinentGroupings worksheet). OMIWNG presence is further positively correlated with the 2000-2010 period (0.24) and negatively correlated with the 2011-2023 period (-0.16; see the DateGroupings worksheet). Interestingly, OMIWNG exhibits a negative correlation (-0.12) with animal isolates and a slight positive correlation with human isolates (0.06; see the HostGrouping worksheet).

In contrast, the OMIUNG variant appeared in only 6% of RefSeq genomes, mainly in *CC30* and *CC121* isolates, and with a negative correlation to *CC5* (-0.16) and *CC8* (-0.14). OMIUNG was found to be slightly positively correlated with strains derived from Europe (0.04) but negatively correlated with those from North America (-0.10). Additionally, the OMIUNG variant exhibited a small negative correlation with human hosts (-0.14).

### Associations exist between Clonal Complex and enterotoxin gene composition

Pearson correlation coefficients were also calculated between Clonal Complex status and enterotoxin gene presence/absence more broadly, with all values to be found in the RefSeqCC worksheet of Supplementary File 2 and selected examples shown in Table 1. For example, both constituents of the *sek-seq* gene cluster showed a strong positive correlation with *CC8* strains. While the same enterotoxin genes exhibited a weaker negative correlation with *CC5, CC5* had a strong positive correlation with *sed-ser-selj-sep*, another commonly observed gene cluster.

**Table 1:** Selected enterotoxin genes and their Pearson correlation coefficient values with membership of CC5 and CC8.

| Enterotoxin gene | CC5 | CC8 |
| --- | --- | --- |
| *sek* | -0.27 | 0.61 |
| *seq* | -0.27 | 0.62<sup>332</sup> |
| *sed* | 0.43 | -0.10<sup>333</sup> |
| *ser* | 0.42 | -0.10 |
| *selj* | 0.42 | -0.11<sup>334</sup> |
| *sep* | 0.31 | -0.12<sup>335</sup> |

Correlation coefficients between Sequence Types and enterotoxin genes were found typically to mirror the values of their associated Clonal Complex to the same enterotoxin genes, as seen in the RefSeqST worksheet of Supplementary File 2. For example, *ST8* displayed the strongest negative correlation with the egc enterotoxin gene cluster genes (e.g. between -0.37 and -0.43 for the OMIWNG genes) and a strong positive correlation with *sek* and *seq* (0.54), mirroring the associations observed for *CC8* isolates. Infrequently, Sequence Types will exhibit relationships with enterotoxin genes that are not found in their respective Clonal Complex, such as for Sequence Type *ST609. ST609* possesses a moderately high correlation coefficient with *seb* (0.26) which is not mirrored in the parent Clonal Complex, *CC8* (0.03), suggesting a more complex relationship with certain lineages in this case.

### Associations exist between enterotoxin gene clusters and genes

Pearson correlation coefficients between enterotoxin gene and gene cluster presence within isolates also varied considerably. Table 2 highlights some of the larger values identified. For example, possession of an *sek*-*seq* gene cluster was suggested by the value of -0.43 to be antagonistic to the OMIWNG variant of the egc enterotoxin gene cluster. Notably, *sek*-*seq* is rare in isolates belonging to certain Clonal Complexes such as CC5, CC22, CC30, CC45 and CC121, where the OMIWNG variant is relatively common (see RefSeqCC worksheet of Supplementary File 2).

**Table 2:**
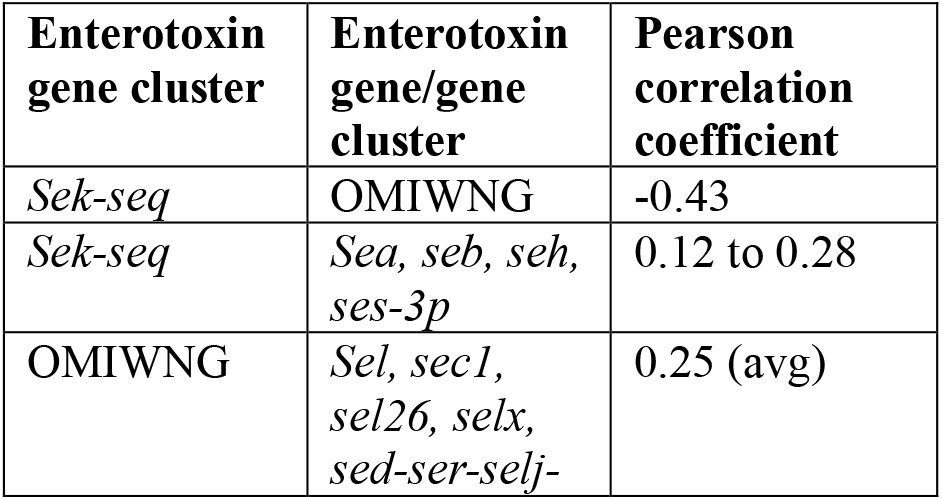
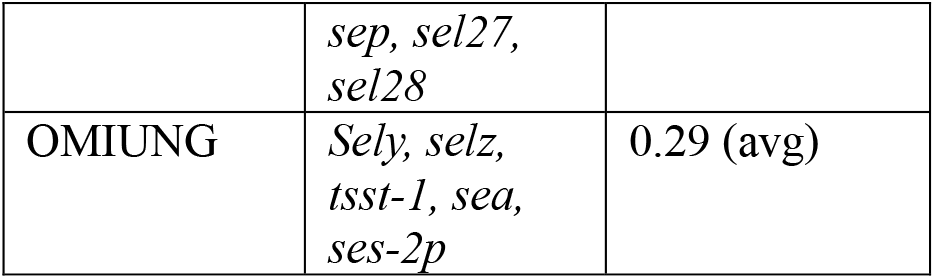
Enterotoxin genes and enterotoxin gene clusters exhibit a broad range of correlation coefficient values with other gene clusters or with individual enterotoxin genes.

Enterotoxin gene groupings underpinned via gene cluster composition or co-occurrence on mobile genetic elements can sometimes be inferred using the correlation and dataset presence values. Such groupings easily evident from these datasets include *sed-ser-selj, sek-seq, sec3-tsst-1, sel-sec1/sec2/sec3, ses-set* and *sel31-sel32*, whereas other associations such as *sel26*-*selx* likely reflect their shared chromosomal location, relatively rare amongst the gene family members. Enterotoxin genes not included in these gene groupings may still be associated with them but more loosely. For example, *seb* presents a positive correlation with *sek-seq* at 0.28, though not as strong as the correlation between *sek* and *seq* at 1.0 (see the RefSeqEnterotoxins worksheet in Supplementary File 2). These results are likely the consequence of the presence of *sek* and *seq* on pathogenicity islands (e.g. SaPI1, 3 and 5) and prophages but with the addition of *seb* on only some of these elements (e.g. SaPI3).

### Mining isolate metadata reveals further patterns of enterotoxin gene association

#### Geographical distribution

Of the 15,887 *S. aureus* isolates analysed, 13,775 originated from 83 countries and 2,112 from unknown locations. Grouped by continent, the data included 5,453 samples from North America, 3,878 from Europe, 2,986 from Asia, 515 from Africa, 551 from South America and 392 from Oceania as shown in Table 3. North America exhibited the highest Pearson correlation coefficient with the egc gene cluster (OMIWNG) at 0.16 with 59% of isolates sourced from this region containing the gene cluster (see the ContinentGroupings worksheet in Supplementary File 2). In contrast isolates derived from Africa and Asia showed the lowest egc gene cluster prevalence at 13%. The *sed-ser-selj* enterotoxin gene cluster was most strongly associated with North America with a correlation coefficient of 0.26, while *seb* was most strongly correlated with Asia at 0.15. Furthermore, the *tsst-1* gene was found to be weakly negatively associated with North America with a value of -0.1.

**Table 3:** Pearson correlation coefficients between the source continents of the isolates in the RefSeq dataset to source host and source time-period. The isolate counts for each of the source metadata categories are displayed.

| Continent | Animal | Human | Unknown | Isolates | Pre-2000s | 2000-2010 | 2011-2023 | Unknown |
| --- | --- | --- | --- | --- | --- | --- | --- | --- |
| North America | -0.157 | 0.312 | -0.232 | **5,453** | -0.042 | 0.256 | -0.167 | -0.083 |
| South America | -0.033 | 0.117 | -0.105 | **551** | -0.001 | -0.034 | 0.081 | -0.064 |
| Europe | 0.121 | -0.164 | -0.096 | **3,878** | 0.109 | 0.212 | -0.080 | -0.194 |
| Asia | 0.116 | -0.043 | -0.031 | **2,986** | -0.046 | -0.252 | 0.407 | -0.207 |
| Oceania | 0.030 | 0.068 | -0.093 | **392** | 0.035 | -0.040 | 0.082 | -0.070 |
| Africa | 0.097 | -0.030 | -0.032 | **515** | -0.024 | -0.080 | 0.144 | 0.820 |
| Unknown | -0.114 | -0.258 | 0.035 | **2,112** | -0.029 | -0.257 | -0.291 | 0.709 |
| **Isolates** | **1,571** | **9,857** | **4,459** | **15,887** | **273** | **5,233** | **7,576** | **2,805** |

Clonal Complex distribution similarly varied geographically. *CC5* isolates were found to be dominant in North and South America at 52% and 49% of all isolates respectively and negatively correlated with Europe (-0.23), contained in only 10% of isolates. Interestingly, *CC22* exhibited the opposite trend (e.g.Pearson’s r=0.352 and with a prevalence of 28% in Europe). *CC93* isolates were largely confined to Oceania and *CC1* displayed a moderate positive correlation with Asia (0.175).

#### Temporal trends

Isolation dates spanned from 1884 to 2023 and were grouped here into four time periods: pre-2000s (273 isolates), 2000-2010 (5,233 isolates), 2011-2023 (7,576 isolates) and Unknown (2,805 isolates). The constituent genes within the egc gene cluster were most prevalent within the 2000-2010 group, being found in 64% of isolates and with a strong preference for the OMIWNG variant (see the DateGroupings worksheet in Supplementary File 2). Prevalence declined post-2011 at 40% with a relative increase in OMIUNG gene cluster variants. The *sed-ser-selj* gene cluster followed a similar temporal pattern to the egc gene cluster. *Seb* was more common in pre-2000s isolates (23%), while *tsst-1* increased in frequency over time (from 4% to 7% between pre-2000s and 2011-2023).

Temporal shifts in Clonal Complex distribution were also observed. *CC8* was predominant pre-2000s (46%) but declined thereafter, while *CC5* increased during 2000-2010 (37%). Post-2011, unknown CCs became more common (25%), potentially indicating broader collection of isolates. Most early isolates (pre-2000s) were from Europe (165/276), whereas recent isolates (2011-2023) while still predominantly originating from Europe and North America now showed greater inclusion from Asia, Africa, South America and Oceania, likely underpinning the increased lineage diversity observed.

#### Host Association

Human-derived isolates correlated positively with *CC5* and *CC8* (0.26 and 0.13) and negatively with *CC22* (-0.23; see the HostGrouping worksheet in Supplementary File 2). Animal-derived isolates were primarily associated with unknown CCs (0.23) and negatively correlated with *CC5* (-0.16). Isolates from unknown hosts further showed a modest positive correlation coefficient with *CC22* (0.30). Geographical differences in host distributions were also evident (see the ContinentGroupings worksheet in Supplementary File 2). South America had the highest proportion of human-derived isolates (92%), Europe showed the most negative correlation with human origin (-0.16) with a high proportion of animal and unknown host origins. Animal-host isolates increased over time, from 3% pre-2000s to 15% in 2011-2023. Isolates with unknown collection dates were predominantly of human or unknown host origin.

### HDBSCAN clustering of RefSeq strains suggests complex relationships between bacterial lineages and enterotoxin gene composition

The HDBSCAN analysis of the RefSeq dataset generated 45 clusters and one noise cluster which was further sub-clustered into 15 sub-clusters according to the scheme presented in the Methods, with the HDBSCAN cluster denoted for each strain within the RefSeqDataset worksheet in Supplementary File 2. The 21 clusters containing all strains harbouring an egc enterotoxin gene cluster are referred to as EGCClusters and the remaining 24 clusters (excepting the noise cluster) are referred to as NonEGCClusters. Section 6 of Supplementary File 1 shows the preferences of each enterotoxin gene for the EGCCluster NonEGCCluster and Noise cluster HDBSCAN cluster types. For example, the *selz* gene shows a high preference for EGCClusters, whereas *sek* and *seq* are mainly found within NonEGCCluster strains. While a few genes other than the chromosomally located *selx* and *sel26* show a high preference for two distinct cluster types, only *sea, seb* and *sely* are found frequently in both EGCCluster and NonEGCCluster strains.

Figure 4 outlines the composition of each Clonal Complex according to these two HDBSCAN cluster types. Distinct HDBSCAN egc presence / absence patterns can be observed for all Clonal Complexes barring *CC1*, which contains a mixture of both EGCCluster and NonEGCCluster strains. All Clonal Complexes contained at least three isolates found in the noise cluster.

#### EGCClusters

The 21 EGCClusters, comprised of 15 OMIWNG- and 6 OMIUNG-associated groups, encompassed 7,178 *S. aureus* isolates (6,360 OMIWNG and 818 OMIUNG). These clusters contained 235 distinct Sequence Types, averaging 16 STs per cluster. Collectively, OMIUNG clusters mainly represented strains from two Clonal Complexes: four clusters comprised *CC30* isolates, one *CC121* and one a mix of *CC121* and one or more unknown Clonal Complexes (see the HDBSCANLineage worksheet in Supplementary File 2). OMIWNG clusters were dominated by isolates from *CC5* (in 11 of 15 clusters), *CC1, CC22* and *CC45. CC22* had the highest proportion of isolates within the EGCClusters at 97.5%, followed by *CC45* (87.6%) and *CC121* (84.9%). *CC5* was the most prevalent Clonal Complex overall, accounting for 51% of all EGCCluster isolates.

The largest EGCCluster contained 2,466 isolates (cluster 43) and featured the OMIWNG enterotoxins genes with *sel26* and *selx*, with 61% of isolates from *CC5*. Clusters 0, 35, 42 and 44 contained variations of the *sed-ser-selj-sep* enterotoxin group, mostly dominated by *CC5* isolates, with cluster zero presenting a combination of *CC5* and *CC45* isolates. Cluster 0 also included both *ses-set* and *ser-selj-sep* gene groups, with membership largely restricted to *CC45* isolates.

#### NonEGCClusters

The 24 NonEGCClusters comprised 7,734 *S. aureus* isolates, spanning 414 unique Sequence Types (average of 25 STs per cluster). Strains belonging to five Clonal Complexes were prominent: *CC1, CC8, CC15, CC93* and *CC97. CC8* was the most abundant (3,378 isolates), present in 14 clusters, while unknown Clonal Complexes appeared in 21 clusters, totalling 2,603 isolates. The largest cluster, cluster 29, (2,821 isolates) contained only the *sel26* and *selx* enterotoxin genes and is considered the “default” cluster. This cluster included eight known Clonal Complexes, though strains without designated Clonal Complexes were more prevalent. Of the strains harbouring the *sek-seq* enterotoxin gene pair, 95% were found in the NonEGCClusters, often also containing the *sea* and *seb* enterotoxin genes (thereby forming a co-located cluster of *sek, seq, sea* and *seb*) across eight HDBSCAN clusters. Cluster 24 (906 isolates) lacked any enterotoxin genes and included all Clonal Complexes except *CC93*, which was exclusively associated (excepting 3 strains within the noise cluster) with cluster 25 that contained *sel26, selx* and *selz*.

#### Noise Cluster

Finally, the noise cluster comprised 975 *S. aureus* isolates, 6.1% of the RefSeq dataset. *CC5* accounted for 385 isolates, the most of any Clonal Complex, followed by the Unknown Clonal Complexes at 238 isolates. Of the 667 Sequence Types in the complete dataset, 123 were represented in the noise cluster, including 51 Sequence Types which only appear in the noise cluster. All enterotoxin genes appeared at least once in the noise cluster. Unsurprisingly *sel26* and *selx* were the most populous enterotoxin genes, followed by the components of the OMIWNG egc enterotoxin gene cluster. Four enterotoxin genes were found only within the noise cluster: *ses-2p, sel31, sel32* and *sel33*. Another fourteen enterotoxin genes or gene variants (*sec2, sec3, see, seh, sel, ses, ses-3p, set, selv, selz, sel29p, sel30, sel34* and *sel35)* in addition to *tsst-1* have 20% or higher presence in the noise cluster despite accounting for only 6.1% of isolates in the RefSeq dataset.

### Association rule mining exhibits connections between isolate enterotoxins, lineages and metadata

Association rule mining unearthed weighted connections between enterotoxin genes, lineages and metadata. Figures 5, 6 and Figure S2 in Supplementary File 2 are network representations of the mined rules, including weighted edges displaying connection strength and arrows indicating the dominant direction of a connection (though note connections may be either unidirectional or bidirectional). The network of rules shown in Figure 5 displays the relationships between enterotoxin genes. For example, the constituents of the enterotoxin gene grouping *sep, ser, selj* and *sed* – known to be often co-located due to their presence on various plasmids - display a strong relationship between themselves, as does the strongest displayed group of enterotoxin genes: *sek, seq, seh, sea* and *ses-3p*. Members of this latter grouping also exhibit strong external relationships. For example, *seb* has strong relationships with both *sek* and *seq*, both of which also connect to *sely*. Figure 5 also shows that the egc gene cluster variant OMIWNG connects strongly to the *ser-sep-selj-sed* grouping, while OMIUNG and OMIWNG share mostly weaker relationships with *tsst-1* and *sely*.

**Figure 5:**
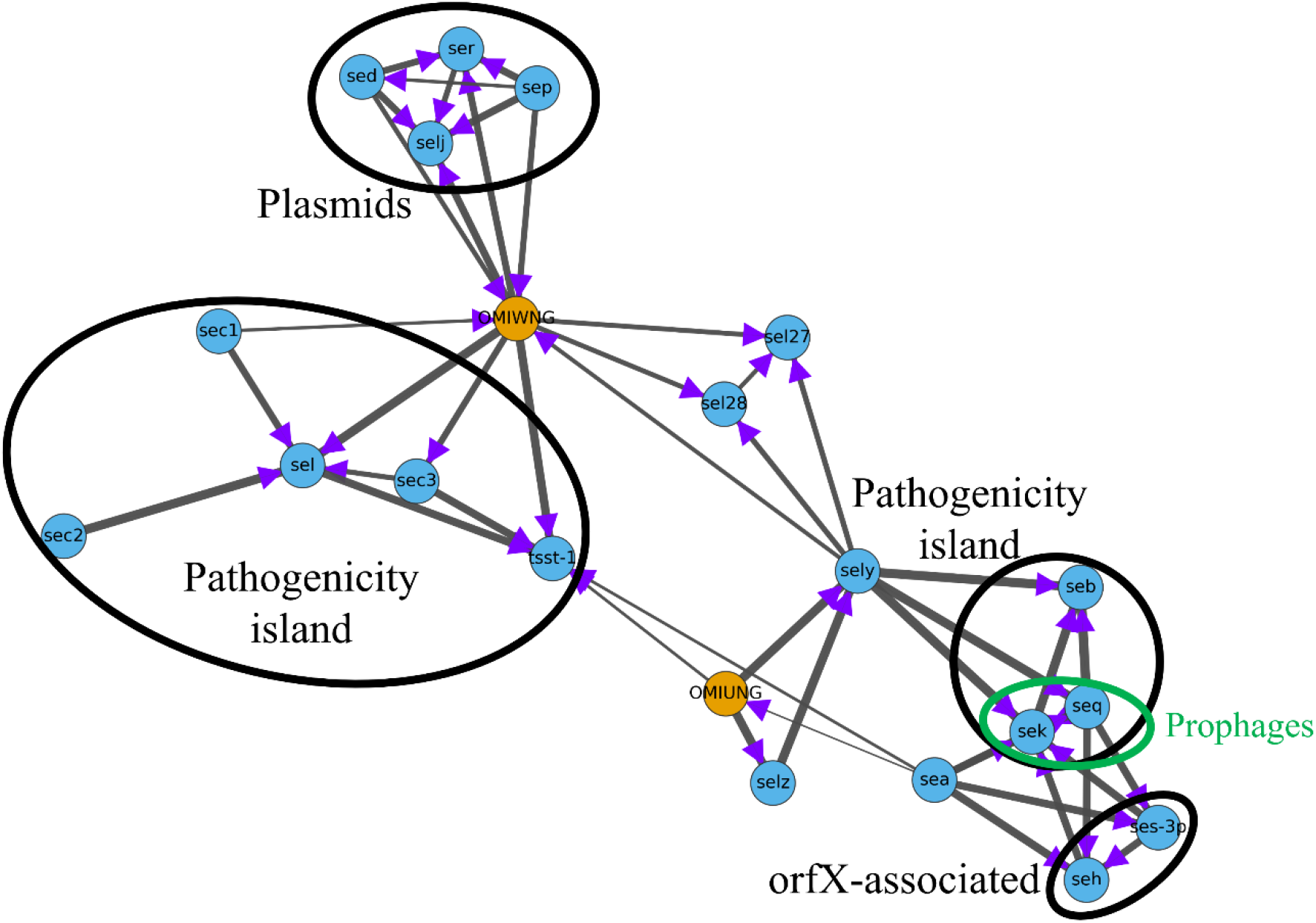
Network of Apriori association rule mined enterotoxin gene relationships. The edge width defines the strength of relationship between enterotoxin genes. The egc gene cluster enterotoxin genes were refined to their related egc cluster variant and *sel26* and *selx* were removed due to their prevalence across the dataset. The enterotoxin genes not found in the figure were not referenced in any association rules and therefore were excluded from this network. Purple arrows indicate the stronger of the two potential associations represented by each edge (e.g. possession of the *sec2* enterotoxin gene strongly suggests the co-presence of the *sel* gene due to the thick edge between the two vertices representing these genes and the arrow pointing towards *sel*). The previously known co-location of enterotoxin genes due to their association with pathogenicity islands, plasmids, prophages and other mobile genetic elements (e.g. see Argudin et al., 10) is indicated by ovals superimposed onto relevant sections of the graph, indicating the potential biological significance of these computationally mined graphical structures.

**Figure 6:**
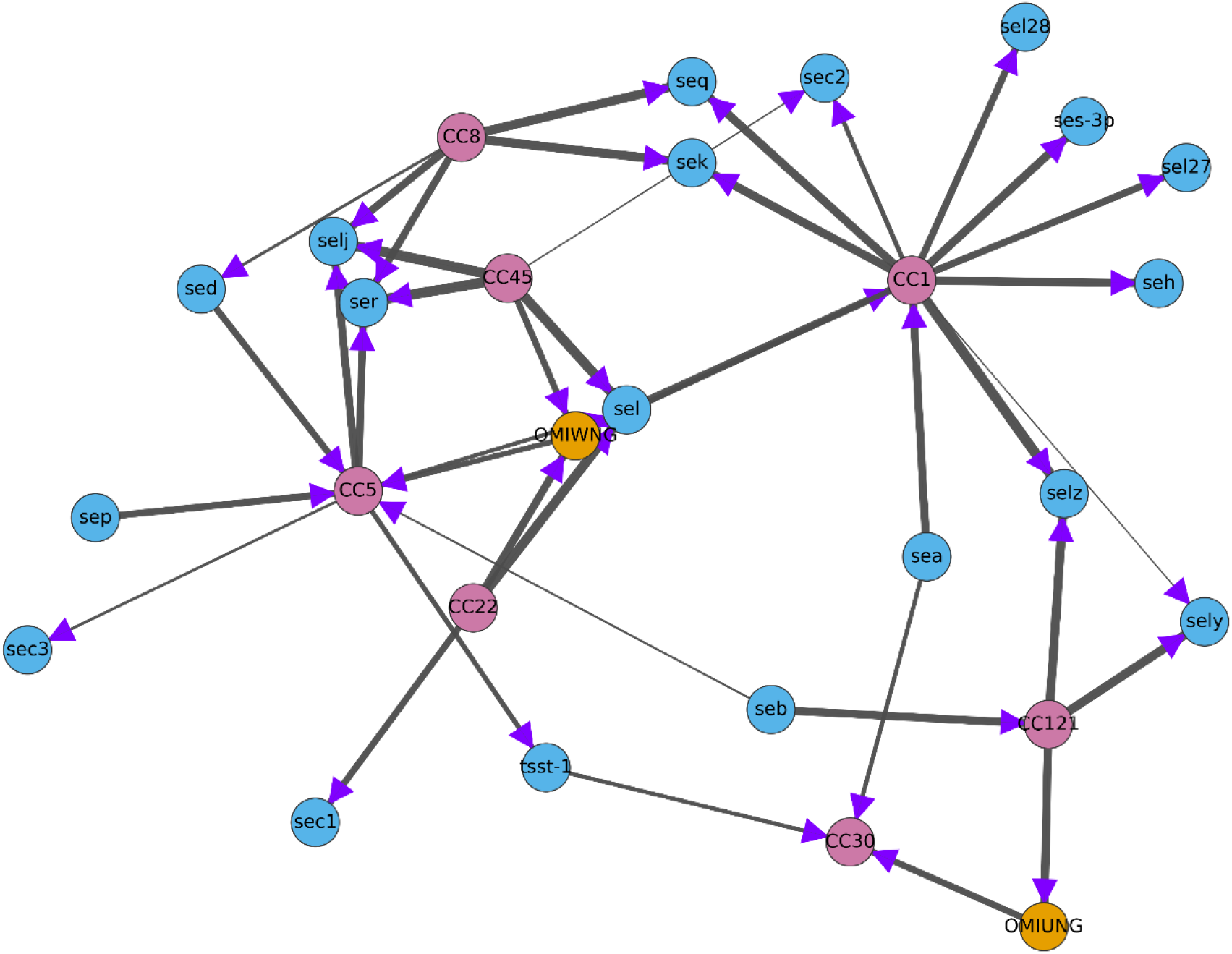
A Clonal Complex-enterotoxin gene network was constructed using the *apriori* association rule mining algorithm, with edge weights calculated using the method described in Sections 2 and 3 of Supplementary File 1. Enterotoxin gene associations were only included if the relationship between two enterotoxin genes exceeded the minimum threshold defined by the algorithm. Purple arrows indicate the direction of the stronger of the two associations between any pair of nodes, again only displayed if the relationship exceeds the minimum threshold. To prevent biased outcomes, *sel26* and *selx* were removed and the egc enterotoxin gene cluster was abstracted into the two main variants OMIUNG and OMIWNG.

The network of enterotoxin genes, gene cluster variants and Clonal Complexes shown in Figure 6 further reflects many of the results previously outlined. For example, OMIUNG is seen to be strongly associated with *CC121* and *CC30*, though with differing directions of association, whereas OMIWNG is related to *CC1, CC5, CC22* and *CC45*. Amongst the enterotoxin genes and gene clusters, OMIWNG exhibits the highest number of connections to Clonal Complexes at 4 connections, followed by *sel, ser* and *selj* at 3, indicating a broad lineage spread of these sub-structures. Seven enterotoxin genes (plus *tsst-1*) have 2 Clonal Complex associations and a further 8 have one only, suggesting a more limited ‘niche’ inheritance of these genes. Of the seven Clonal Complexes shown here, *CC1* exhibits the most connections with 12 – likely a reflection of its prominence across different HDBSCAN cluster types - followed by *CC5* with 8 and *CC8* with 6.

## 8. Discussion

In this study, we investigated enterotoxin gene presence, lineage information and isolate metadata across the genome assemblies of 15,887 *Staphylococcus aureus* isolates. While many previous studies of pathogenic bacteria have typically analysed fewer than 100 strains (e.g. (34,35)), our dataset has provided a much broader view of strains collected and sequenced over the past century. The dataset encompassed an extensive set of enterotoxin gene profiles, detailed MLST-based lineage classifications and metadata including country of origin, year of isolation and host organism. This expansive approach provided a robust overview of trends in *S. aureus* genome sequencing and may prove valuable for understanding bacterial behaviour and guiding future outbreak prevention strategies. Furthermore, by applying machine learning techniques such as clustering, correlation analysis and association rule mining to this dataset, we were able to shed light on the high genetic diversity within the RefSeq *S. aureus* collection and reveal notable associations between the analysed factors across isolates from different backgrounds.

The identification of two novel enterotoxin-like genes, *sel34* and *sel35*, expands the total number of known enterotoxin genes to 35, (18 confirmed “se” and 17 “sel” forms, plus 5 variants). Both genes were detected in isolates from East Africa, with phylogenetic analyses placing *sel34* near *seb* and *sel35* within the *sea/sel26* clade. Notably *seb*, along with the toxic shock associated superantigen *tsst-1*, may be particularly significant to the scientific community given its potential to induce immune system collapse (34,37,38), so determining the function of a closely related gene may be of interest. The egc enterotoxin gene cluster also plays a critical role in *S. aureus* epidemiology, influencing colonization rates, pathogenicity, and disease outcomes. It has been suggested that the presence of egc gene cluster superantigens is negatively correlated with the severity of sepsis, such that while non-egc superantigens are secreted predominantly during the late stationary phase, egc-related superantigens are released during exponential growth before ceasing at higher bacterial densities (7). Other studies have linked egc gene cluster presence with reduced infection severity, providing a protective effect on the bacterium, while being more commonly found within airway isolates (8,39,40). In our study, egc enterotoxin genes were present in 49% of the RefSeq dataset and typically appeared in two variants, OMIUNG and OMIWNG, with the latter dominating (43% of all isolates) compared to OMIUNG (6%). Our use of MLST-based lineage information enabled us to identify dominant lineages of isolates sequenced to date, with over half of sequenced strains found to be within Clonal Complexes CC5 and CC8. However, it is currently unknown whether these high counts are a result of biased sampling or the global dominance of these lineages. Notably, isolates from *CC121* carried the highest number of enterotoxin genes with 11 on average, whereas those from *CC15* and *CC97* carried as few as two. Unsurprisingly, Clonal Complexes harbouring eight or more enterotoxins were strongly associated with the egc enterotoxin gene cluster, which comprises six genes for most strains, while strains lacking this gene cluster exhibited significantly lower median enterotoxin gene counts.

Strains belonging to particular Clonal Complexes often exhibited affinities for specific enterotoxin genes or groups of enterotoxin genes and vice versa . For example, the OMIWNG variant of the egc gene cluster was predominantly found in *CC5, CC22* and *CC45* strains, whereas *CC30* and *CC121* were associated with the OMIUNG variant. Interestingly, the *sed-ser-selj+sep* gene group correlated strongly with *CC5* but negatively with *CC22*, both OMIWNG-related complexes, showing that associations were often not straightforward. Additionally, undefined or unknown Clonal Complexes, which account for 20% of the RefSeq dataset, showed a strong negative correlation with the egc gene cluster. Given that the egc gene cluster is reportedly present in 50–60% of *S. aureus* isolates, the underrepresentation in these undefined lineages may skew the observed proportions of egc-related versus non-egc complexes and should be considered when drawing conclusions about *S. aureus* feature prevalence.

HDBSCAN clustering analysis yielded 44 clusters and one noise cluster. Roughly a half of these groups were associated with egc enterotoxin gene cluster presence, encompassing 7,178 isolates and representing 235 unique Sequence Types. As anticipated, clusters linked to the OMIWNG variant were predominantly composed of Clonal Complexes *CC1, CC5, CC22*, and *CC45*, reinforcing our earlier findings from correlation analysis. In contrast, the 24 clusters largely lacking egc gene cluster presence included 7,734 isolates and 414 unique Sequence Types, displaying a markedly broader lineage composition. Notably, *CC8* was the most populous lineage, suggesting that *CC8* is the primary Clonal Complex for *S. aureus* isolates which lack an egc gene cluster. The noise cluster, representing 6% of the RefSeq dataset (975 isolates), exhibited considerable diversity in enterotoxin gene composition, with all enterotoxin genes represented at least once and some (e.g., *ses-3p, sel33, sel31*, and *sel32*) unique to this cluster. The disproportionate presence of certain enterotoxin genes in the noise cluster may reflect small-scale variations that arise from the relatively low minimum threshold used in the HDBSCAN algorithm.

The apriori mined rules between enterotoxin genes, lineages and metadata strongly supported the earlier correlation and cluster analyses in this study. The networks demonstrate both connections and strength of relationship between various factors of *S. aureus* enterotoxin-associated epidemiology. Given the very large quantities of association data produced by the correlation and clustering analyses, this type of machine learning method may prove highly useful going forward in capturing the strongest signals and displaying them visually. They could be embedded in bacterial surveillance dashboards, for example. Within this study, the networks provide support for egc gene cluster associations to specific Clonal Complexes, metadata and other enterotoxin genes that have also been found through the correlation and HDBSCAN analyses. Crucially, sub-structures of networks involving enterotoxin genes were shown to map onto enterotoxin-associated mobile genetic elements, demonstrating the link to the underlying biological system is strong.

The integration of GenBank metadata with the RefSeq dataset enabled a comprehensive exploration of *S. aureus* isolate information based on source location, collection year, and host organism. The RefSeq isolates originated from 84 countries, encompassed 48 host types, and spanned 139 years. Figure 7 presents the data in a mirrored stacked stream graph, with colour gradients in the graph revealing the temporal distribution of isolate collection and sequencing. North America led in the total number of isolates whose genomes were sequenced, with North America and Europe showing high isolate counts during 2000-2010 (2,704 and 1,957, respectively), whereas most sequenced isolates from other continents were collected between 2011 and 2023. The largest temporal increase in isolate collection occurred in Asia, possibly reflecting scientific expansion in countries such as Japan and China, with 90% of Asian isolates originating from 2011-2023 (41). A growth in the sequencing of isolates sourced from animals in Asia and Europe is also seen, particularly since 2011, potentially a consequence of the increasing prominence of One Health approaches within bacteriology.

**Figure 7:**
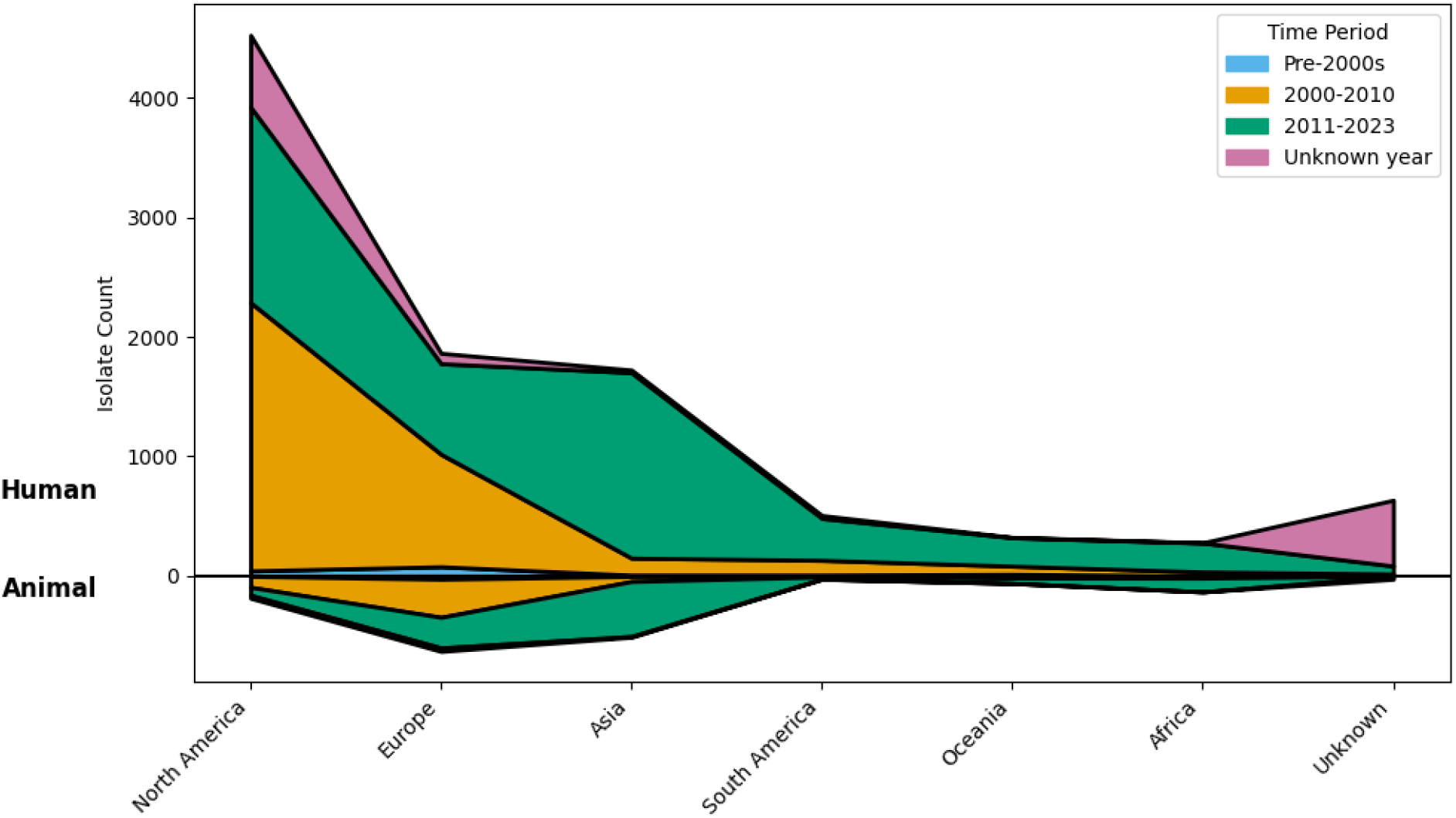
A mirrored stacked stream graph generated using all aspects of isolate metadata within the RefSeq dataset. The isolate count extends out from the mirror line, above the line depicting a human host or below an animal host. The colours of the stream stack depict isolate collection year with each stream increment representing a continent (shown along the x-axis of the figure).

There are, however, some limitations to our study. Notably, lineage typing tools and databases inevitably lag behind lineages emerging in the scientific literature (35,42,43). Consequently, approximately 20% of the isolates (3,178) lacked an assigned Clonal Complex. Furthermore, between 13-26% of isolates lacked useful metadata, meaning that while they could be examined for enterotoxin gene composition and lineage assignment, this data could not be associated to information on collection host, year or location. Despite these limitations, our study identified two new members of the growing enterotoxin gene family, notably from animal sources. We also showed that machine learning methods can be used to uncover plausible information about the relationships between gene family members and other features of the isolates undergoing analysis. In future, it would be interestng to more closely align this information with genomic structures such as plasmids, pathogenicity islands and prophages, which could provide information on emerging genetic structures underpinning the proliferation of this important gene family.

## 1.5 Abbreviations

CC: Clonal complex
ST: Sequence Type
MLST: Muli-Locus Sequence Typing
MRSA: Methicillin-resistant *Staphylococcus aureus*

## 9. Author statements

### 9.1 Author contributions

Contributions have been attributed by the CRediT system as follows.

Conceptualization: V.M., R.L., J.D. Formal Analysis: A.U. Methodology: A.U., V.M., R.L., J.D. Writing – original draft: A.U. Writing – review & editing: V.M., R.L., J.D.

## 9.4 Acknowledgements

We would like to thank all those who made their datasets publicly available, thereby making possible studies such as these, and all those who look after the data once submitted.

