## Supplementary file 1 for "Machine learning approaches for the identification and analysis of enterotoxin genes in *Staphylococcus aureus* genomes"

### Supplementary File 1 for "Machine learning approaches for the identification and analysis of enterotoxin genes in *Staphylococcus aureus* genomes" by Uttin *et al.*, 2026

#### Section 1 – HDBSCAN cluster labelling with t-SNE

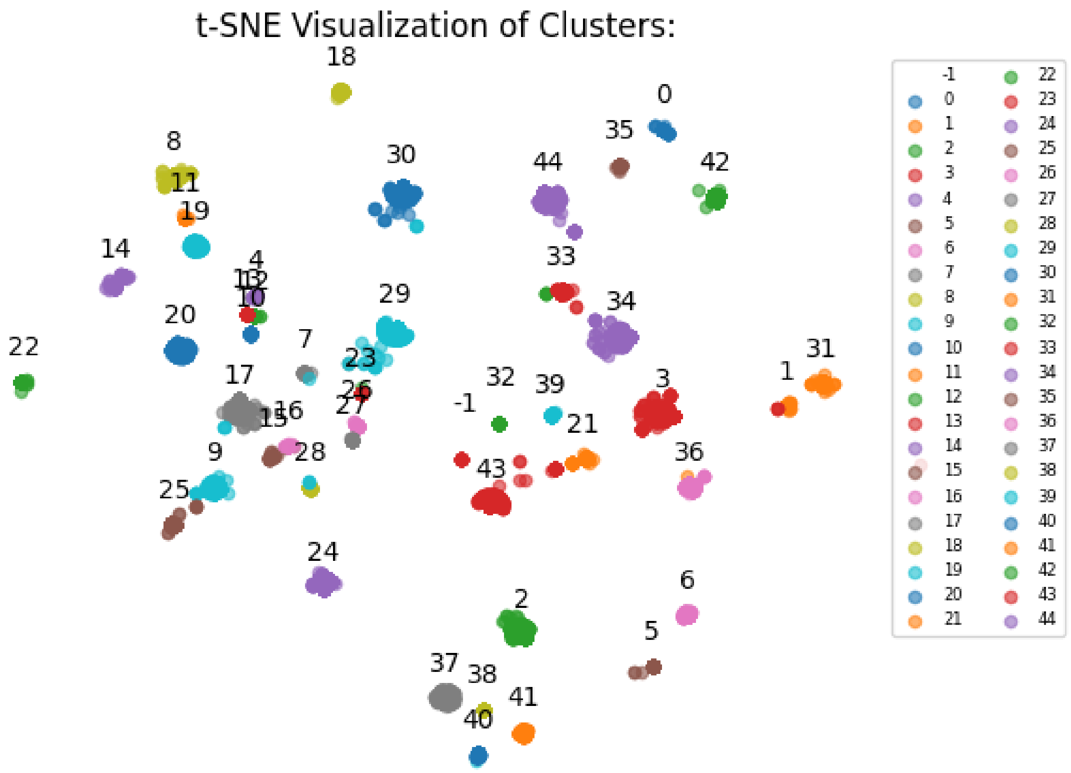


***Figure S1*: HDBSCAN isolate cluster labels superimposed on the selected t-SNE representation of the RefSeq dataset enterotoxin gene presence/absence data.** t-SNE scaled the RefSeq's 15,887 isolates using the 39 enterotoxin features to a two-dimensional positional value displayed above. The cluster labels generated by HDBSCAN were applied to the positional values by colouring the points. The t-SNE model used the parameters: *Components: 2, Perplexity: 100, Iter: 2000* and *Learning rate: 200.* The HDBSCAN model used the parameters: *Minimum cluster size: 40, Minimum sample size: 20, Epsilon value: 0.3, Metric: manhattan* and *Cluster selection metric: eom.*

#### Section 2 - Association rule mining to network flattening

Consider two hypothetical rules:

- Antecedents: *sely* and *sel33*, Consequents: *selj*, Confidence: 0.9
- Antecedents: *selj* and *sely*, Consequents: *sel33* and *ses*, Confidence: 0.8

By directionally flattening these rules, we create a dataset that lists each antecedent followed by its consequents and their averaged confidence values. For these two rules, our dataset would appear as:

| **Enterotoxin** | **Associated Entity** | **Confidence Calculation** | **Confidence** |
|---|---|---|---|
| sely | sel33 | (0.9 + 0.8) / 2 | 0.85 |
| sely | selj | (0.9 + 0.8) / 2 | 0.85 |
| sely | ses | 0.8 | 0.8 |
| sel33 | sely | 0.9 | 0.9 |
| sel33 | selj | 0.9 | 0.9 |
| selj | sely | 0.8 | 0.8 |
| selj | sel33 | 0.8 | 0.8 |
| selj | ses | 0.8 | 0.8 |
| ses | (No associations) | | |

On completion of the flattening, edge strength must be determined before edge weight can be calculated. Edge strength is determined by averaging the confidence scores of the two nodes connected by that edge. If the flattening has produced a unidirectional relationship between two nodes, then the averaging is not required.

Examples using the dataset generated above:

- sely <-> selj, strength: (((0.9 + 0.8) / 2) + 0.8 ) / 2

- selj <-> ses, strength: 0.8

Edge weight is subsequently determined by conducting min-max normalisation across the newly calculated edge strengths. This produces a dataset of nodes connected by weighted edges, visualised by network graphs. This unidirectional flattening of association rules to edge weighting process remains consistent when expanding to include CC and strain source information.

### Section 3 - Formula Summary for Association Rule Mining

#### Define input sets and elements:

Let:

- $X = \{X_0, X_1, \dots, X_n\}$ be the set of antecedents.
- $Y = \{Y_0, Y_1, \dots, Y_n\}$ be the set of consequents.
- $p$ denotes a rule, with $Z_p$ representing the strength of rule $p$.

#### Define dictionary structure:

- Let $D$ be the dictionary where each key $X_i \in X$ has a value that is another dictionary $D_{X_i}$
- $D_{X_i}$ has keys $Y_j \in Y$ with corresponding values as a list $\left[Z_{X_i,Y_j}, c_{X_i,Y_j}\right]$ $where$:
  - $Z_{X_i,Y_j}$ is the cumulative strength of rules associating $X_i$ and $Y_j$
  - $c_{X_i,Y_j}$ is the count of rules associating $X_i$ and $Y_j$

#### Update rules and calculations:

- For each rule $p$ involving $X_i \to Y_j$ with strength $Z_p$:
  - If $Y_j$ is not in $D_{X_i}$, initialise $D_{X_i}\left[Y_j\right] = \left[Z_p, 1\right]$.
  - If $Y_j$ is already in $D_{X_i}$, update it as follows:
    - $D_{X_i}\left[Y_j\right][0] = D_{X_i}\left[Y_j\right][0] + Z_p$
    - $D_{X_i}\left[Y_j\right][1] = D_{X_i}\left[Y_j\right][1] + 1$

#### Calculating average strengths:

- At the end, for each $X_i$ and $Y_j$ pair, compute the average strength:

$$\text{Strength}_{X_i,Y_j} = \frac{D_{X_i}\left[Y_j\right][0]}{D_{X_i}\left[Y_j\right][1]}$$

#### Calculate combined strength for each pair $X_i$ and $X_j$:

- For each pair of antecedents $X_i$ and $X_j$ (where $i \neq j$), calculate the combined strength:

$$\text{Combined Strength}_{X_i,X_j} = \frac{\text{Strength}_{X_i,Y_{X_j}} + \text{Strength}_{X_j,Y_{X_i}}}{2}$$

- Here, $\text{Strength}_{X_i,Y_{X_j}}$ is retrieved from $D_{X_i}\left[Y_{X_j}\right]$ and similarly for $\text{Strength}_{X_j,Y_{X_i}}$

#### Min-Max normalisation for combined strength values:

- Define $S_{\min}$ and $S_{\max}$ as the minimum and maximum values among all Combined $\text{Strengths}_{X_i,X_j}$ in the list.
- For each $\text{Combined Strength}_{X_i,X_j}$, calculate the normalised strength:

$$\text{Normalised Strength}_{X_i,X_j} = \frac{\text{Combined Strength}_{X_i,X_j} - S_{\min}}{S_{\max} - S_{\min}}$$

#### Final list structure with normalised combined strengths:

- The final result is a list of triplets for each pair $\left(X_i, X_j\right)$:

$$\left[X_i, X_j, \text{Normalised Strength}_{X_i,X_j}\right]$$

- Which will look like:

$$\left[X_0, X_1, \text{Normalised Strength}_{X_0,X_1}\right], \left[X_0, X_2, \text{Normalised Strength}_{X_0,X_2}\right], \dots, \left[X_i, X_j, \text{Normalised Strength}_{X_i,X_j}\right]$$

#### Section 4 - Association rule mined networks

**a)**

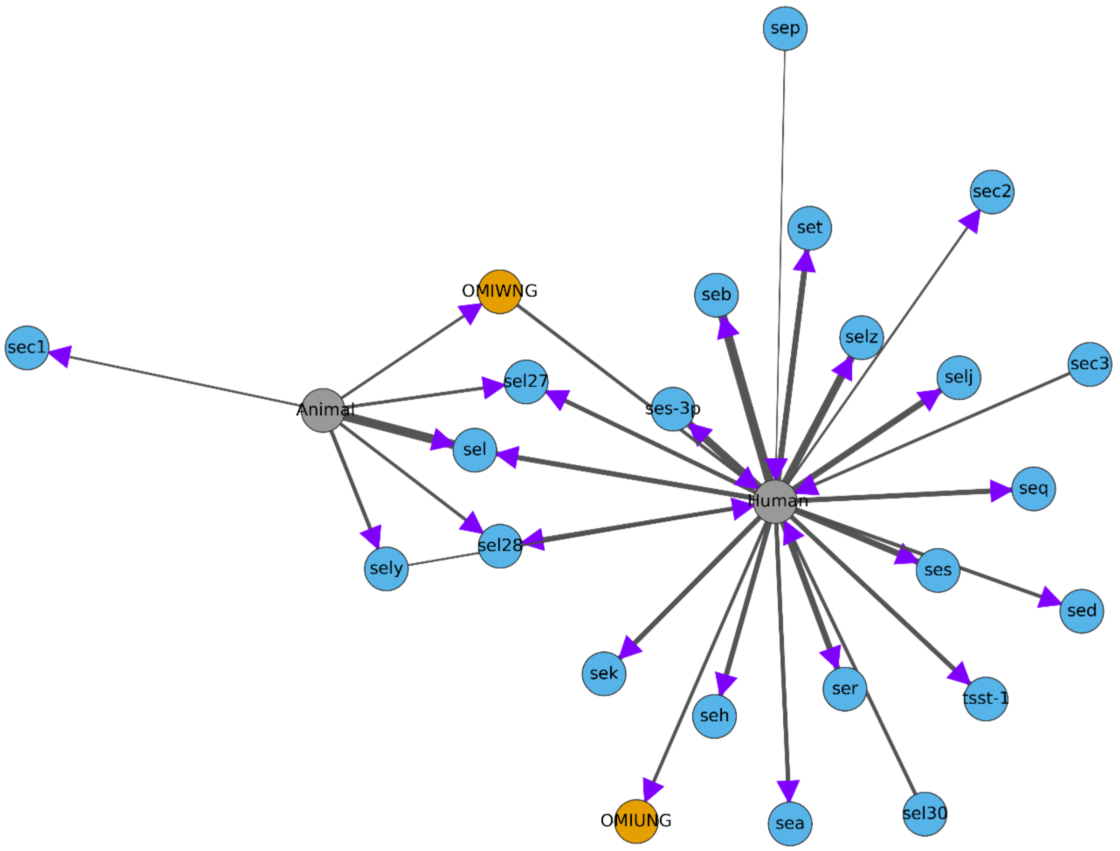


**b)**

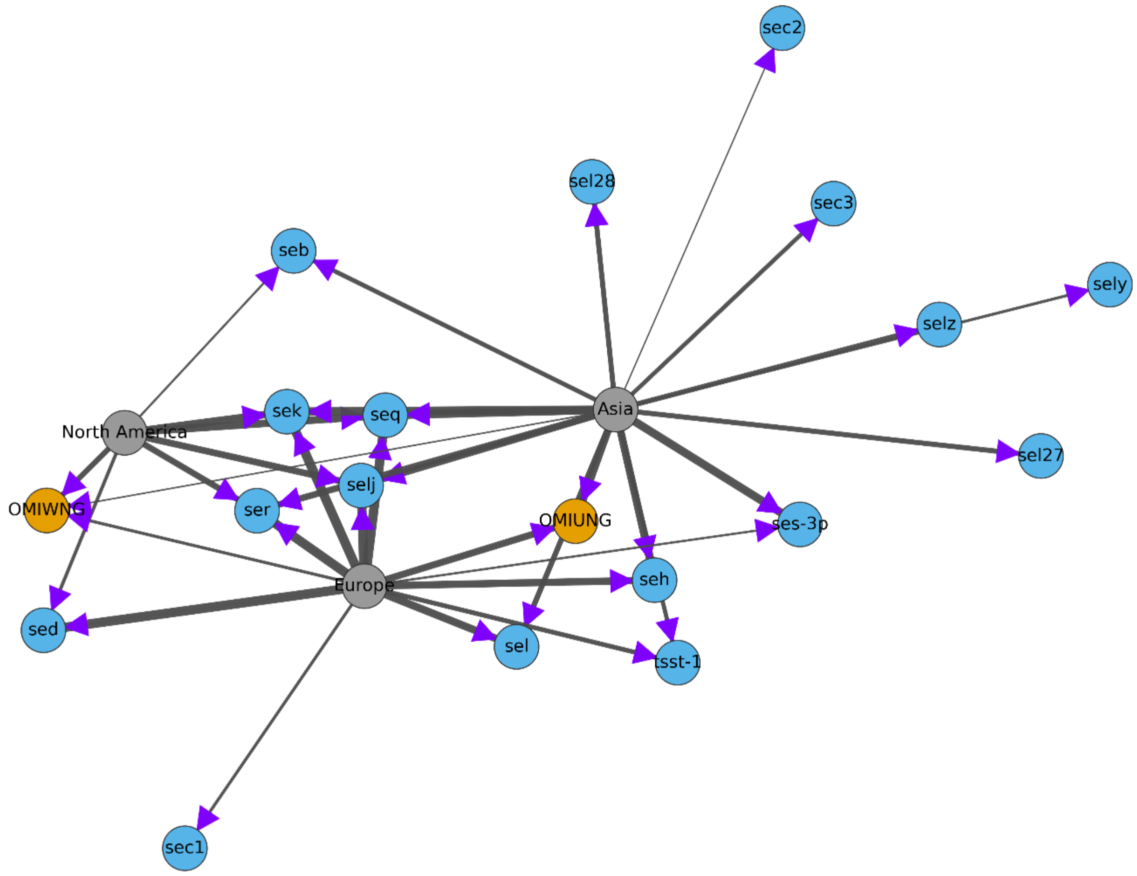

**c)**

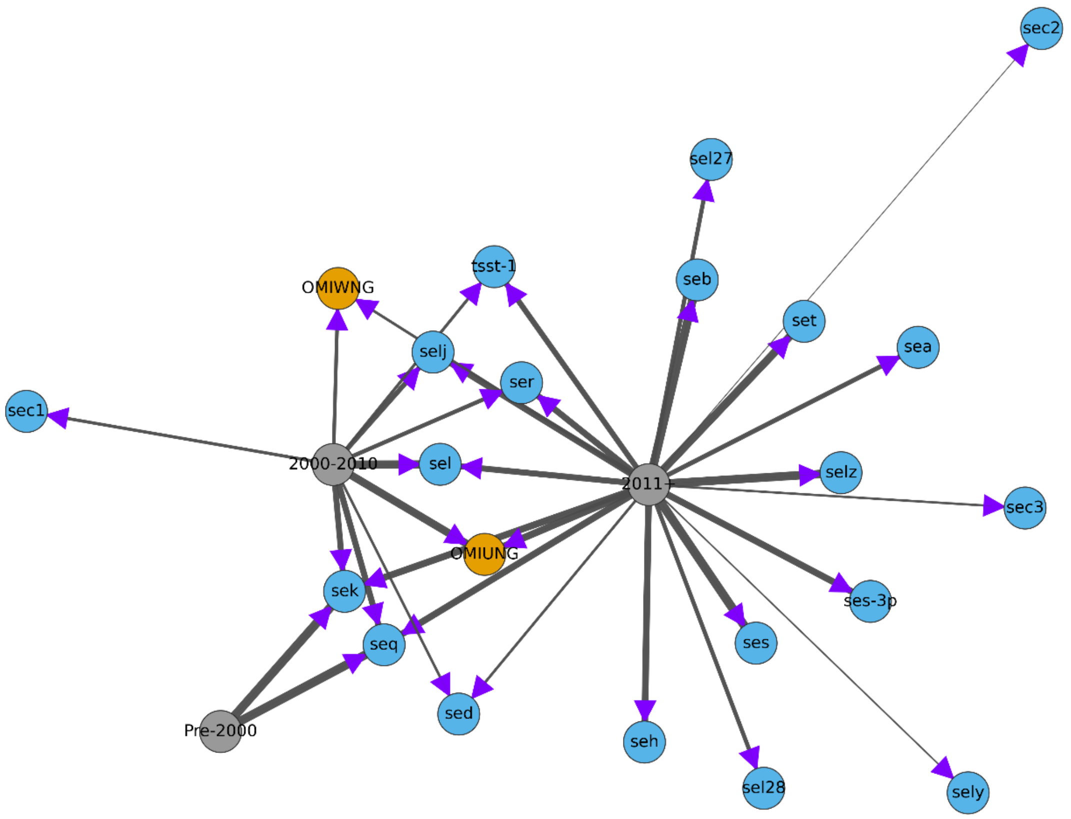


**Figure S2: Apriori association rule mining network between *S. aureus* enterotoxin genes and strain metadata, specifically a) host, b) continent and c) time period.** Edge weights were calculated using the method described in Sections 2 and 3 above. Relationships were only included if the relationship between the enterotoxin genes and the metadata exceeded the minimum threshold as defined by the algorithm. Purple arrows indicate the direction of the stronger of the two associations between any pair of nodes, again only displayed if the relationship exceeds the minimum threshold. The *sel26* and *selx* enterotoxin genes were removed due to their high prevalence and the egc gene cluster was consolidated into the two main variants, OMIWNG and OMIUNG.

#### Section 5 - Novel enterotoxin gene sequences identified in this study

***sel34***

ATGAGAAAAAACTTTAAAAAACTAATTAAAAACATTTGTGTTGCATCTCTAGCTGTTTTAA
CCATACTTATTATAGGTGCAGATAGAGCTGAAGCACAACCAGATCCTGAGCCAGGGGAATT
GCATAAATCTAGTCAATTTACAGGTTTAATGGAAAACCTAAGAGTTTTATATGACGATTATC
ATGTAGAAGCCAGTAATGTTAGCACTGTTGAGCAATTTTTGAATTTTGATTTAATTTTTCCA
ATTGTTGACCATCAATTACATAATTATGATAAAGTGAGAATAGAGTTTGCTGATAAACTATTA
GCAGATAAGTTTAAAGGTAAAAATGTAGATGTATTTGGAACAAATTATTATAGACATTGTTA
TTTCTCAGAAGAAGATAAGAATGAAGAAAATGAAACAAATGGAGCTAAAAATAAAACATG
CATGTATGGAGGAGTAACAAAACATGATGATAATCATTATGATTCAAGTGATGAACCCTCG
AATGCAAAAAATGTATTAGTACAAGTGTTTATAGATAAAGTCAATGTCATGTCATTTGACAT
TCAAACTAATAAAAAACAAGTTACTGCTCAAGAACTTGACTATAAAGCAAGAAATTATTTG
ATTAAACATAAAAATTTATATGAATTTAAAAACTCACCTTATGAAACAGGATATATTAAGTTT
ATTGAAAATGGCAATGAATCTTTTTGGTACGATTTGATGCCCTCTCCAGGTAAAACATTTGA
TCAATTTAGGTATCTTATGATTTATAATGATAACAAAATAGTTGATTCTAATAAGACAAAGAT
ACAAGTGTTTTTAACAAAAAAGTAA

***sel35***

ATGTTTAAAAAGTTTTTTACATTAATTTTAAGTTTGGTATTGCTTTATCCGTTGTTAAGCAAT
TTTTGTAATGCTAATGCTGAGAAAATTCCAAAACCTGAGGATTTACACAAGAAGAGTGAA
GTGGAACATTTAAATAATTTAAGGCATGCCTATTCATTTGGTAATTTTACAATAGCCAATGAT
AAAAAAACGGATGAACAATTTTTATCAAATTCTTTGCTTTTTTATGGTTTCTTTACTGATCA
CCCTATATATAAAGACCTACTAATATCTTTTGATTCTGACAGTCACGCAAAAAAATTTTTAG
GGAAATCAATAGATATTTATGGGATTGGTTTCGGTAATAATTGTGAAGGGGGGACACCAGG
GAAAACACAATGCATGTATGGCGGTGTTACTCCTCATGAAAACAACATAATGACTAATGAT-
AAAAATATACCTATAAACCTATGGTTGGACGGAAAACAAACAGAAGTAGCTTACTCAACA
GTAACAACCAATAAGAAAATTGTTACTGTTCAAGAATTAGATGCCAAAGTAAGAAAATATT
TAAGTGATAAATATAAAATATATGAAACAGATATATGGGGGGGACGATTCAAAGAGGATTAG
TTCAGTTTGATGGAGCAAGTGAAAAAGTATCGTTTGATTTATATGGAGCAAAAGGTAAATT
TGCAGAATCATTTTTAAAAATTTATAAAGACAATAAAACAATCAGTTCAGAAAACTTACAC
ATTGACATATATTTATATACAAAATAA

#### Section 6 - HDBSCAN Clustering enterotoxin gene preference

***Table S1*: This table outlines the preferences of each enterotoxin gene with the HDBSCAN cluster groups (egc-related or non-egc related clusters).** Each row has been individually normalised with the min-max scores of corresponding integer values of each row. This leaves individual gene preferences without influence from other enterotoxin genes. The averages and deviations have not been normalised.

| Toxin | EGC | NonEGC | Noise |
|---|---|---|---|
| seo | 1 | | 0.09 |
| sem | 1 | | 0.09 |
| sen | 1 | | 0.08 |
| seg | 1 | | 0.08 |
| sei | 1 | | 0.08 |
| selu | 1 | | 0.22 |
| selw | 1 | | 0.07 |
| sely | 1 | 0.47 | |
| selz | 1 | | 0.02 |
| sel | 1 | | 0.21 |
| sec1 | 1 | | 0.02 |
| sec2 | 1 | | 0.56 |
| sec3 | 1 | | 0.71 |
| tsst-1 | 1 | | 0.40 |
| selv | 0.21 | | 1 |
| sel26 | 1 | 0.93 | |
| selx | 0.99 | 1 | |
| sek | | 1 | 0.05 |
| seq | | 1 | 0.05 |
| sea | 0.36 | 1 | |
| seb | 0.28 | 1 | |
| sed | 1 | 0.07 | |
| ser | 1 | 0.04 | |
| selj | 1 | 0.04 | |
| sep | 1 | 0.16 | |
| sel30 | 1 | | 0.62 |
| seh | | 1 | 0.49 |
| seh-2p | | 1 | 0.17 |
| ses-2p | | | 1 |
| ses-3p | | 1 | 0.34 |
| sel27 | 1 | | 0.12 |
| sel28 | 1 | | 0.11 |
| sel29p | | 1 | 0.5 |
| ses | 1 | | 0.39 |
| set | 1 | | 0.32 |
| see | | | 1 |
| sel33 | | | 1 |
| sel31 | | | 1 |
| sel32 | | | 1 |
| sel34 | | 1 | 1 |
| sel35 | | 1 | 1 |
| Average | 0.63 | 0.31 | 0.34 |
| STD | 0.47 | 0.45 | 0.38 |
